# Sparse Tensor Decomposition for Haplotype Assembly of Diploids and Polyploids

**DOI:** 10.1101/130930

**Authors:** Abolfazl Hashemi, Banghua Zhu, Haris Vikalo

## Abstract

A framework that formulates haplotype assembly as sparse tensor decomposition is proposed. The problem is cast as that of decomposing a tensor having special structural constraints and missing a large fraction of its entries into a product of two factors, U and 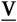; tensor 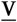 reveals haplotype information while U is a sparse matrix encoding the origin of erroneous sequencing reads. An algorithm, AltHap, which reconstructs haplotypes of either diploid or polyploid organisms by solving this decomposition problem is proposed. Starting from a judiciously selected initial point, AltHap alternates between two optimization tasks to recover U and 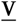 by relying on a modified gradient descent search that exploits salient structural properties of U and 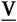. The performance and convergence properties of AltHap are theoretically analyzed and, in doing so, guarantees on the achievable minimum error correction scores and correct phasing rate are established. AltHap was tested in a number of different scenarios and was shown to compare favorably to state-of-the-art methods in applications to haplotype assembly of diploids, and significantly outperform existing techniques when applied to haplotype assembly of polyploids.

## 1 INTRODUCTION

Fast and accurate DNA sequencing has enabled unprecedented studies of genetic variations and their effect on human health and medical treatments. Complete information about variations in an individual’s genome is given by haplotypes, the ordered lists of single nucleotide polymorphisms (SNPs) on the individual’s chromosomes [37]. Haplotype information is of fundamental importance for a wide range of applications. For instance, when the corresponding genes on a homologous pair of chromosomes contain multiple variants, they could exhibit different gene expression patterns. In humans, this may affect an individual’s susceptibility to diseases and response to therapeutic drugs, and hence suggest directions for medical and pharmaceutical research [11]. Haplotype information also enables whole genome association studies that focus on the so-called tag SNPs [19], representative SNPs in a region of the genome characterized by strong correlation between alleles (i.e., by high linkage disequilibrium). Moreover, haplotype sequences can be used to infer recombination patterns and identify genes under positive selection [36]. In addition to the SNPs and minor structural variations found in a healthy individual’s genome, complex chromosomal aberrations such as translocations and nonreciprocal structural changes – including aneuploidy – are present in cancer cells. Cancer haplotype assembly enables identification of “driver” mutations and thus helps to understanding the mechanisms behind the disease and discovery of its genetic signatures.

Haplotype assembly from short reads obtained by high-throughput DNA sequencing requires partitioning (either directly or indirectly) the reads into *K* clusters (*K* = 2 for diploids, *K* = 3 for triploids, etc.), each collecting the reads corresponding to one of the chromosomes. If the reads were free of sequencing errors, this task would be straightforward. However, sequencing is erro-neous – state-of-the-art platforms have error rates on the order of 10^−3^ – 10^−2^. This leads to ambiguities regarding the origin of a read and therefore renders haplotype assembly challenging. For this reason, the vast majority of haplotype assembly techniques attempts to remove the aforementioned ambiguities by either discarding or altering sequencing data; this has led to the minimum fragment removal, minimum SNP removal [26], maximum fragments cut [16], and minimum error correction formulations of the assembly problem [29]. Most of the recent haplotype assembly methods (see, e.g., [7, 25, 31, 32, 40]) focus on the minimum error correction (MEC) formulation where the goal is to find the smallest number of nucleotides in reads that need to be changed so that any read partitioning ambiguities would be resolved. It has been shown that finding optimal solution to the MEC formulation of the haplotype assembly problem is NP-hard [7, 10, 26]. In [39], the authors used a branch-and-bound scheme to minimize the MEC objective over the space of reads; to reduce the search space, they relied on a bound on the objective obtained by a random partition of the reads. Unfortunately, exponential growth of the complexity of this scheme makes it computationally infeasible even for moderate haplotype lengths. Integer linear programming techniques have been applied to haplotype assembly in [9], but the approach there fails at computationally difficult instances of the problem. More recently, fixed parameter tractable (FPT) algorithms with runtimes exponential in the number of variants per read [6, 22] were proposed; these methods are well-suited for short reads but become infeasible for the long ones. A dynamic programming scheme for haplotype assembly of diploids proposed in [21] is also exponential in the length of the longest read. A probabilistic dynamic programming algorithm that optimizes a likelihood function generalizing the MEC objective is developed in [25]; this method is characterized by high accuracy but is significantly slower than the previous heuristics. [31, 32] aim to process long reads by developing algorithms for the exact optimization of weighted variants of the MEC score that scale well with read length but are exponential in the sequencing coverage. These methods, along with ProbHap [25], struggle to remain accurate and practically feasible at high coverages (e.g., higher than 12 [25]).

The computational challenges of optimizing MEC score has motivated several polynomial time heuristics. In a pioneering work [28], a greedy algorithm seeking the most likely haplotypes was used to assemble haplotypes of the first complete diploid individual genome obtained via high-throughput sequencing. To compute posterior joint probabilities of consecutive SNPs, Bayesian methods relying on MCMC and Gibbs sampling schemes were proposed in [4] and [24], respectively; unfortunately, slow convergence of Markov chains that these schemes rely on limits their practical feasibility. Following an observation that haplotype assembly can be interpreted as the clustering problem, a max-cut formulation was proposed in [3]; an efficient algorithm (HapCUT) that solves it and significantly outperforms the method in [28] was developed and has been widely used subsequently. A flow-graph based approach in [1], HapCompass, re-examined fragment removal strategy and demonstrated superior performance over HapCut. Other recent diploid haplotype assembly methods include a greedy max-cut approach in [17], convex optimization program for minimizing the MEC score in [13], and a communication-theoretic interpretation of the problem solved via belief propagation (BP) in [34]. Note that deep sequencing coverage provided by state-of-the-art high-throughput sequencing platforms and the emergence of very long insert sizes in recent technologies (e.g., fosmid [17]) may enable assembly of extremely long haplotype blocks but also impose significant computational burden on the methods above.

Increased affordability, capability to provide deep coverage, and longer sequencing read lengths also enabled studies of genetic variations of polyploid organisms. However, haplotype assembly for polyploid genomes is considerably more challenging than that for diploids; to illustrate this, note that for a polyploid genome with *k* haplotype sequences of length *m*, under the all-heterozygous assumption there are (*k* − 1)^*m*^ different genotypes and at least 2^(*m* − 1)^ (*k* − 1)^*m*^ different haplotype phasings. In part for this reason relatively fewer methods for solving the haplotype assembly problems in polyploids have been developed. In fact, with the exception of HapCompass [1], SDhaP [13] and BP [34], the above listed methods are restricted to diploid genomes. Other recent techniques capable of reconstructing haplotypes for both diploid and polyploid genomes include HapTree [5], a Bayesian method to find the maximum likelihood haplotype shown to be superior to HapCompass and SDhaP (see, e.g., [30] for a detailed comparison), and H-PoP [40], the state-of-the-art dynamic programming method that significantly outperforms the schemes developed in [1, 5, 13] in terms of accuracy, memory consumption, and speed.

In this paper, we propose a unified framework for haplotype assembly of diploid and polyploid genomes based on sparse tensor decomposition; the framework essentially solves a relaxed version of the NP-hard MEC formulation of the haplotype assembly problem. In particular, read fragments are organized in a sparse binary tensor which can be thought of as being obtained by multiplying a matrix that contains information about the origin of erroneous sequencing reads and a tensor that contains haplotype information of an organism. The problem then is recast as that of decomposing a tensor having special structural constraints and missing a large fraction of its entries. Based on a modified gradient descent method and after unfolding the observed and haplotype information bearing tensors, an iterative procedure for finding the decomposition is proposed. The algorithm exploits underlying structural properties of the factors to perform decomposition at a low computational cost. In addition, we analyze the performance and convergence properties of the proposed algorithm and determine bounds on the minimum error correction (MEC) scores and correct phasing rate (CPR) – also referred to as reconstruction rate – that the algorithm achieves for a given sequencing coverage and data error rate. To the best of our knowledge, this is the first polynomial time approximation algorithm for haplotype assembly of diploids and polyploids having explicit theoretical guarantees for its achievable MEC score and CPR. The proposed algorithm, referred to as AltHap, is tested in applications to haplotype assembly for both diploid and polyploid genomes (synthetic and real data) and compared with several state-of-the-art methods. Our extensive experiments reveal that AltHap outperforms the competing techniques in terms of accuracy, running time, or both. It should be noted that while state-of-the-art haplotype assembly methods for polyploids assume haplotypes may only have biallelic sites, AltHap is capable of reconstructing polyallelic haplotypes which are common in many plants and some animals, are of particular importance for applications such as crop cultivation [35], and may help in reconstruction of viral quasispecies [33]. Moreover, unlike several state-of-the-art haplotype assembly methods that have complexity which scales exponentially with either read length (e.g., [9]) or coverage (e.g., [25]), AltHap’s iterative steps are linear in both; this makes AltHap well-suited for haplotype assembly from long sequencing reads and deep coverage data. Indeed, we confirm this claim by performing extensive experiment using simulated datasets.

## 2 MATHEMATICAL MODEL AND PROBLEM FORMULATION

We briefly summarize notation used in the paper. Bold capital letters refer to matrices and bold lowercase letters represent vectors. Tensors are denoted by underlined bold capital letters, e.g., 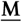. M_::1_ and 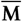 denote the frontal slice and the mode-1 unfolding of a third-order tensor 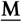, respectively. For a positive integer *n*, [*n*] denotes the set {1 …, *n*}. The condition number of rank-*k* matrix M is defined as *κ* = *σ*_1_/*σ*_*k*_ where *σ*_1_ ≥ · · · ≥ *σ*_*k*_ > 0 are singular values of M. SVD_*k*_ (M) denotes the rank *k* approximation (compact SVD) of M computed by power iteration method [2, 27].

Let 𝓗 = {**h**_1_, …, **h**_*k*_} denote the set of haplotype sequences of a *k*-ploid organism, and let **R** be an *n* × *m* SNP fragment matrix where *n* denotes the number of sequencing reads and *m* is the length of haplotype sequences. **R** is an incomplete matrix that can be thought of as being obtained by sampling, with errors, matrix **M** that consists of *n* rows; each row of **M** is a sequence randomly selected from among *k* haplotype sequences. Since each SNP is one of four possible nucleotides, we use the al-phabet 𝒜 = {(1, 0, 0, 0), (0, 1, 0, 0), (0, 0, 1, 0), (0, 0, 0, 1)} to describe the information in the haplotype sequences; the mapping between nucleotides and alphabet components follows arbitrary convention.

The reads can now be organized into an *n* × *m* × 4 SNP fragment tensor which we denote by 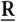. The (*i*, *j*,:) fiber of 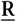, i.e., a one-dimensional slice obtained by fixing the first and second indices of the tensor, represents the value of the *j*^th^ SNP in the *i*^th^ read. Let Ω denote the set of informative fibers of 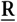, i.e., the set of (*i*, *j*,:) such that the *i*^th^ read covers the *j*^th^ SNP. Define an operator 𝒫_Ω_(.) as

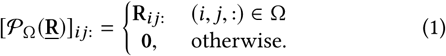

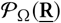 is a tensor obtained by sampling, with errors, tensor 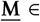 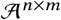 having *n* copies of *k* encoded haplotype sequences as its horizontal slices. More specifically, we can write 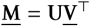, where 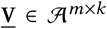 contains haplotype information, i.e., the *j*^th^ vertical slice of 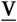, V_:*j*:_, is the encoded sequence of the *j*^th^ haplotype, and U∈ {0, 1}^*n*×*k*^ is a matrix that assigns each of *n* horizontal slices of 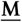 to one of *k* haplotype sequences, i.e., the *i*^th^ row of U, **u**_*i*_, is an indicator of the origin of the *i*^th^ read. Let Φ = {**e**_1_, …, **e**_*k*_}, where e_*l*_ ∈ ℝ^*k*^ is the *l*^th^ standard basis vector having 1 in the *l*^th^ position and 0 elsewhere. The rows of U are standard unit basis vectors in ℝ^*k*^, i.e., **u**_*i*_ ∈ Φ, ∀*i* ∈ [*n*]. This representation is illustrated in Fig. 1 where the (1, 1,:) fiber of 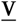 specified with dashed lines is mapped to the (1, 1,:) fiber of 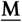 which in turn implies that in the example described in Fig. 1 we have **u**_1_ = **e**_1_.

**Fig. 1.**
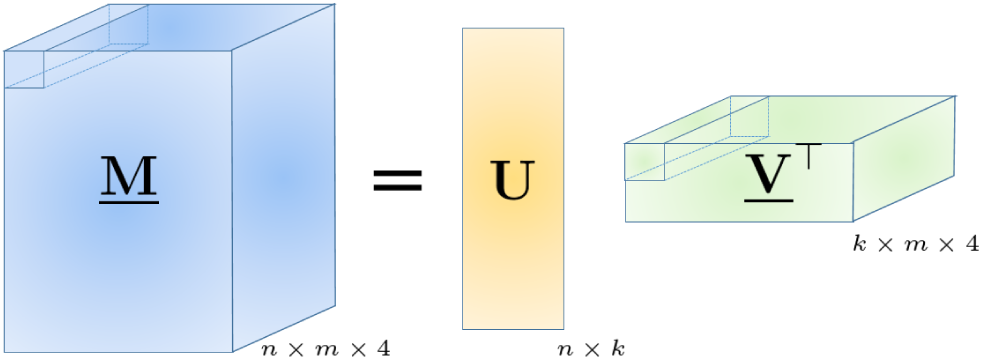
Representing haplotype sequences and sequencing reads using tensors. Tensor 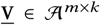 contains haplotype information while matrix U ∈ {0, 1}^*n*×*k*^ assigns each of the *n* horizontal slices of 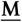 to one of the *k* haplotype sequences, i.e., the *i*^th^ row of U is an indicator of the origin of the *i*^th^ read.

DNA sequencing is erroneous and hence we assume a model where the informative fibers in 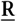 are perturbed versions of the corresponding fibers in 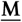 with data error rate *p*_*e*_, i.e., if the (*i*, *j*,:) ∈ Ω fiber in 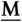 takes value e_*l*_ ∈ 𝒜, R*_ij_*: with probability 1 − *p_e_* equals e_*l*_ and with probability *p*_*e*_ takes one of the other three possibilities. Thus, the observed SNP fragment tensor can be modeled as 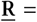 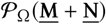 where 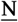 is an additive noise tensor defined as

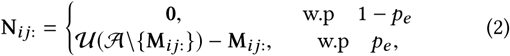

where the notation 𝒰(𝒜\{M_*i j*:_}) denotes uniform selection of a vector from 𝒜\{M_*i j*:_}. The goal of haplotype assembly can now be formulated as follows: *Given the SNP fragment tensor* 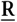*, find the tensor of haplotype sequences 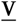 that minimizes the MEC score*.

Next, we formalize the MEC score as well as the correct phasing rate, also known as reconstruction rate, the two metrics that are used to characterize performance of haplotype assembly schemes (see, e.g., [9, 14, 18, 21]). For two alleles **a**_1_, **a**_2_ ∈ 𝒜 ∪ {0}, we define a dissimilarity function *d*(**a**_1_, **a**_2_) as

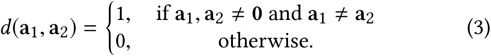

The MEC score is the smallest number of fibers in 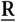 that need to be altered so that the resulting modified data is consistent with the reconstructed haplotype 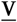, i.e.,

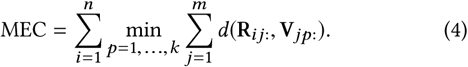

The correct phasing rate (CPR), also referred to as the reconstruction rate, can conveniently be written using the dissimilarity function *d*(., .). Let 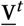 denote the tensor of true haplotype sequences. Then

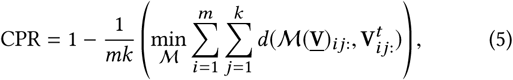

where 𝓜 is a one-to-one mapping from lateral slices of 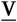 to those of 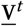, i.e., a one-to-one mapping from the set of reconstructed haplotypes to the set of true haplotypes.

We now describe our proposed relaxation of the MEC formulation of the haplotype asseLmbly problem. Let *p*_*i*_ ∈ [*k*], ∀*i* ∈ [*n*] be defined as *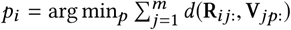*. Notice that for any *j* such that 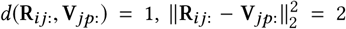. Therefore, by denoting Ω_*i*_ the set of informative fibers for the *i*^th^ read we obtain^1^

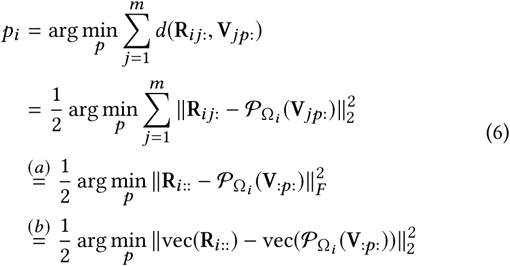

where (*a*) follows from the definition of the Frobenius norm and vec(.) in (*b*) denotes the vectorization of its argument. Let **e**_*p*_ be the *p*^th^ standard unit vector ∀*p* ∈ [*k*]. It is straightforward to observe that the last equality in (6) can equivalently be written as 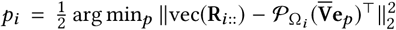 where 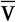 is the mode-1 unfolding of the tensor 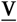. Hence, 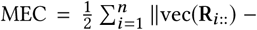 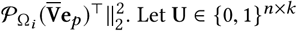 be the matrix such that for its *i*^th^ row it holds that **u**_*i*_ = e_*pi*_. In addition, notice that vec (**R**_*i*::_) is the *i*^th^ row of 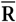. Therefore, from the definition of the Frobenius norm and the fact that 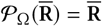 we obtain

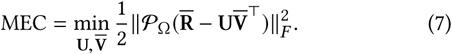

The optimization problem in (7) is NP-hard since the entries of 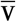 are binary and the objective function is non-convex. Relaxing the binary constraint to 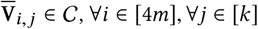, where 𝒞 = [0, 1], results in the following relaxation of the MEC formulation,

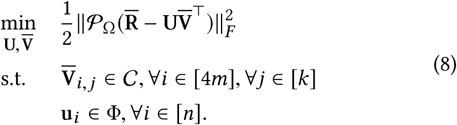

The new formulation can be summarized as follows. We start by finding the so-called mode-1 unfolding of tensors 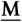 and 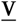 and denote the decomposition 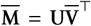, as illustrated in Fig. 2. As implied by the figure, after unfolding, the entries of the (1, 1,:) fiber are mapped to four blocks of 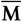 and 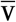 that correspond to the frontal slices of tensors 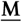 and 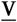, respectively. Then, to determine the haplotype sequence that minimizes the MEC score, one needs to solve (8) and find the optimal tensor decomposition.

**Fig. 2.**
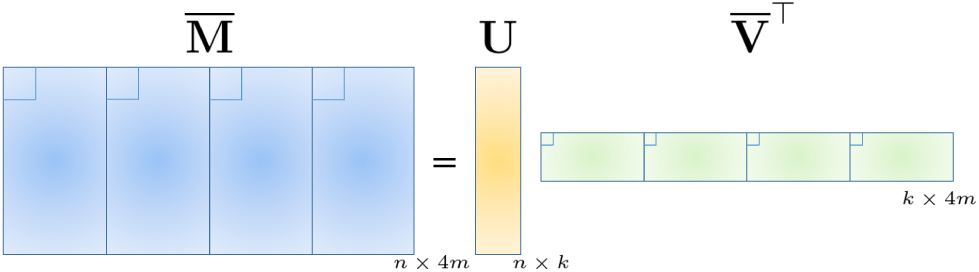
Representing haplotype sequences and sequencing reads using unfolded tensors. Matrix 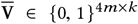 contains haplotype information while matrix U ∈ {0, 1}^*n*×*k*^ assigns each of the *n* rows of 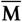 to one of the *k* haplotype sequences, i.e., the *i*^th^ row of U is an indicator of the origin of the *i*^th^ read.

## 3 STRUCTURED TENSOR DECOMPOSITION ALGORITHM

Although the objective function 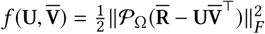 in (8) is convex in each of the factors when the other factor is fixed, 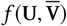 is generally nonconvex. To facilitate computationally efficient search for the solution of (8), we rely on a modified gradient search algorithm which exploits the special structures of **U** and 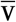 and iteratively updates the estimates 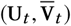 starting from some initial point 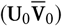. More specifically, given the current estimates 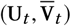, the update rules are

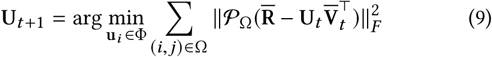

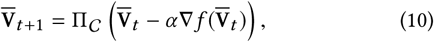

where 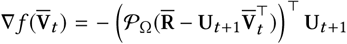 denotes the partial derivative of 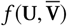 evaluated at 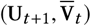, *α* is a judiciously chosen step size, and ∏_𝒞_ denotes the projection operator onto 𝒞. Notice that the optimization in (9) is done by exhaustively searching over *k* vectors in Φ. Since the number of haplotypes *k* is relatively small, the complexity of the exhaustive search (9) is low. The proposed scheme is formalized as Algorithm 1. MATLAB and Python implementations of AltHap are freely available from https://sourceforge.net/projects/althap/.

### Algorithm 1

Structured Tensor Decomposition Algorithm

**Input:** SNP fragment matrix **R**, step size *α*, maximum number of iterations *T*

**Output: 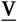**, an estimate of the true haplotype tensor 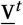

**Preprocessing:** Encode **R** to binary tensor 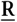 and find the mode-1 unfolding, 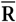

**Initialization:** Compute 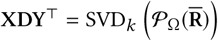 and let 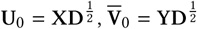. Define Φ = {**e**_1_, …, **e**_*k*_}

**for** *t* = 0; 1; 2; 3 …, *T* − 1 **do**

1. 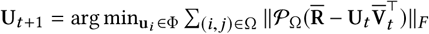

2. 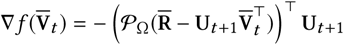

3. 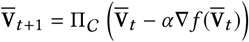

**end for**

Decode 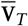 to obtain 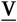

## 4 CONVERGENCE ANALYSIS OF ALTHAP

In this section, we analyze the convergence properties of AltHap and provide performance guarantees in different scenarios.

In the appendix we show that, a judicious choice of the step size *α* according to

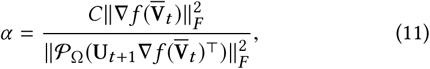

where 𝒞 ∈ (0, 2) is a constant, guarantees that the value of the objective function in (8) decreases as one alternates between (9) and (10), which in turn implies that AltHap converges. The key observation that leads to this result is that 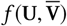 is a convex function in each of the factor matrices and that 𝒞 = [0, 1] is a convex set; hence the projection ∏_𝒞_ in (10) leads to a reduction of 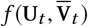 in each iteration *t*.

It is important however to determine the conditions under which the stationary point of AltHap coincides with the global optima of (8). To this end, we first provide the definition of incoherence of matrices [8].

### Definition 4.1.

*A rank-k matrix* M ∈ ℝ^*n*×*m*^ *with singular value decomposition 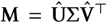 is incoherent with parameter* 1 ≤ *μ* ≤ 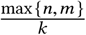 *if for every* 1 ≤ *i* ≤ *n,* 1 ≤ *j* ≤ *m*

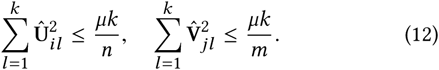

Let each fiber in 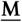 be observed uniformly with probably *p*. Let 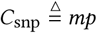 denote the expected number of SNPs covered by each read, and 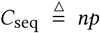 denote the expected coverage for each of the haplotype sequences. Theorem 4.2 built upon the results of [20, 23, 38] states that with an adequate number of covered SNPs, the solution found by AltHap reconstructs 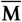 up to an error term that stems from the existence of errors in sequencing reads.

### THEOREM 4.2.

*Assume 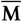 is μ-incoherent. Suppose the condition number of* 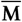 *is κ. Then there exist numerical constants C*_0_, *C*_1_ > 0 *such that if* Ω *is uniformly generated at random and*

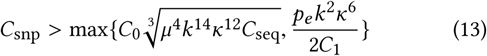

*with probability at least* 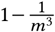 *, the solution 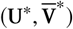 found by AltHap satisfies*

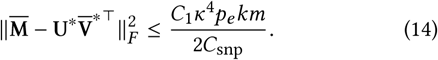

The proof of Theorem 4.2 which is omitted for brevity relies on a coupled perturbation analysis to establish a certain type of local convexity of the objective function around the global optima. Thus, under (13) there is no other stationary point around the global optima and hence, starting from a good initial point, AltHap converges globally. We employ the initialization procedure suggested by [38] – summarized in the initialization step of Algorithm 1 – which is based on a low cost singular value decomposition of 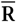 using power method [2, 27] and with high probability lies in the described convexity region of *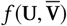*.

*Remark 1:* Under the assumption of 4.2, the Condition 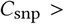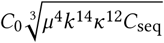 specifies a lower bound on the expected number of covered SNPs, *C*_snp_, that is required for the exact recovery of 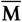 in the idealistic error-free scenario, i.e., for *p*_*e*_ = 0. With higher sequencing coverage, more SNPs are covered by the reads and hence *C*_snp_ required for accurate haplotype assembly scales with *C*_seq_ along with other parameters. Moreover, the term 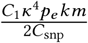 on the right hand side of (14) is the bound on the error of the solution generated by AltHap which increases with the sequencing error rate *p*_*e*_ and ploidy *k*, and decreases with *C*_snp_ and the number of reads *n*, as expected.

*Remark 2:* If 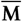 is well-conditioned, i.e., 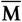 is characterized by a small incoherence parameter *μ* and a small condition number *κ*, the recovery becomes easier; this is reflected in less strict sufficient condition (13s) and improved achievable performance (14). In fact, as we verified in our simulation studies, by using the proposed framework for haplotype assembly, the parameters *μ* and *κ* associated with 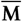 are close to 1 (the ideal case). Theorem 4.3 provides theoretical bounds on the expected MEC scores and CPR achieved by AltHap. (The proof is in the appendix.)

### THEOREM 4.3.

*Under the conditions of Theorem 4.2, with probability at least 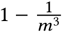 it holds that*

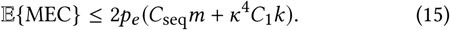

*Moreover, if the reads sample haplotype sequences uniformly, with probability at least* 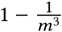 *it holds that*

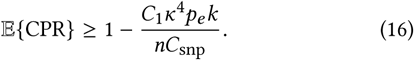

*Remark 3:* The bound established in (15) suggests that the expected MEC increases with the length of the haplotype sequences, sequencing error, number of haplotype sequences, and sequencing coverage. A higher sequencing coverage results in a larger fragment data which in turn leads to higher MEC scores.

*Remark 4:* As intuitively expected, the bound (16) suggests that AltHap’s achievable expected CPR improves with the number of sequencing reads and the SNP coverage; on the other hand, the CPR deteriorates at higher data error rates. Finally, assuming the same sequencing parameters, (16s) implies that reconstruction of polyploid haplotypes is more challenging than that of diploids.

## 5 SIMULATION RESULTS AND DISCUSSION

We evaluated the performance of the proposed method on both experimental and simulated data, as described next. AltHap was implemented in Python and MATLAB, and the simulations were conducted on a single core Intel Xeon E5-2690 v3 (Haswell) with 2.6 GHz and 64 GB DDR4-2133 RAM. The benchmarking algorithms include Belief Propagation (BP) [34], a communication-inspired method capable of performing haplotype assembly of diploid and biallelic polyploid species, HapTree [5], and H-PoP [40], the state-of-the-art dynamic programming algorithm for haplotype assembly of diploid and biallelic polyploid species shown to be superior to HapTree [5], HapCompass [1], and SDhaP [13] in terms of both accuracy and speed [30, 40]. Following the prior works on haplotype assembly (see, e.g., [9, 14, 18, 21]) we use MEC score and CPR to assess the quality of the reconstructed haplotypes.^2^

### 5.1 Experimental data

We first tested performance of AltHap in an application to haplotype reconstruction of a data set from the 1000 Genomes Project – in particular, the sample NA12878 sequenced at high coverage using the 454 sequencing platform. In this work, we take the triophased variant calls from the GATK resource bundle [15] as the true haplotype sequences. We compare the MEC score, CPR, and running time achieved by AltHap to those of H-PoP, BP, and HapTree. All the algorithms used in the benchmarking study were executed with their default settings. The results are given in Table 1. As seen there, among the considered algorithms AltHap achieves the smallest MEC score for nearly all chromosomes and the highest CPR for majority of the chromosomes. Moreover, H-PoP and BP are the fastest and second fastest schemes but their speed comes at the cost of reduced accuracy. On the other hand, AltHap is slightly slower than H-PoP and BP but faster than HapTree.

**Table 1.** Performance comparison of AltHap, H-PoP, BP, and HapTree applied to haplotype reconstruction of the CEU NA12878 data set in the 1000 Genomes Project.

| Chromosome | AltHap |  |  | H-PoP |  |  | BP |  |  | HapTree |  |  |
| --- | --- | --- | --- | --- | --- | --- | --- | --- | --- | --- | --- | --- |
|  | CPR | MEC | t(sec) | CPR | MEC | t(sec) | CPR | MEC | t(sec) | CPR | MEC | t(sec) |
| 1 | 0.974 | **2011** | 11.26 | 0.957 | 2264 | **5.22** | **0.991** | 2321 | 8.17 | 0.841 | 2305 | 15.43 |
| 2 | 0.953 | **2562** | 12.22 | **0.956** | 2971 | **5.65** | 0.895 | 2897 | 9.83 | 0.845 | 2875 | 17.59 |
| 3 | **0.933** | **2084** | 10.38 | 0.912 | 2312 | **6.99** | 0.743 | 2367 | 8.30 | 0.852 | 2363 | 15.06 |
| 4 | 0.969 | **2368** | 12.16 | **0.970** | 2648 | **5.24** | 0.748 | 2613 | 6.76 | 0.835 | 2604 | 18.67 |
| 5 | **0.972** | **1924** | 9.96 | 0.966 | 2103 | **4.67** | 0.882 | 2185 | 4.76 | 0.848 | 2171 | 16.95 |
| 6 | 0.949 | 3687 | 14.17 | **0.952** | **3343** | **4.93** | 0.887 | 3588 | 6.94 | 0.846 | 3583 | 23.86 |
| 7 | **0.970** | **1846** | 11.19 | 0.924 | 1986 | **4.24** | 0.811 | 2073 | 7.88 | 0.847 | 2070 | 13.06 |
| 8 | **0.962** | **1634** | 9.63 | 0.947 | 1848 | **4.14** | 0.885 | 1857 | 8.01 | 0.842 | 1838 | 14.81 |
| 9 | **0.971** | **1272** | 6.42 | 0.910 | 1462 | **3.36** | 0.898 | 1491 | 6.13 | 0.851 | 1479 | 14.90 |
| 10 | **0.968** | **1584** | 7.97 | 0.945 | 1683 | **3.67** | 0.908 | 1839 | 7.18 | 0.857 | 1823 | 12.13 |
| 11 | **0.933** | **1394** | 7.45 | 0.915 | 1553 | **3.71** | 0.756 | 1586 | 6.69 | 0.836 | 1577 | 11.33 |
| 12 | **0.921** | **1423** | 7.12 | 0.903 | 1570 | **3.46** | 0.744 | 1589 | 6.48 | 0.848 | 1589 | 9.97 |
| 13 | **0.970** | **1269** | 4.42 | 0.941 | 1440 | **2.89** | 0.891 | 1409 | 5.38 | 0.828 | 1405 | 9.55 |
| 14 | 0.903 | **857** | 9.53 | **0.971** | 974 | **2.54** | 0.700 | 995 | 4.53 | 0.854 | 987 | 7.79 |
| 15 | 0.972 | **941** | 9.42 | **0.974** | 1039 | **2.40** | 0.746 | 1063 | 3.92 | 0.836 | 1061 | 7.43 |
| 16 | **0.967** | 1198 | 5.40 | 0.935 | **1192** | **2.47** | 0.797 | 1269 | 4.42 | 0.851 | 1273 | 8.13 |
| 17 | **0.975** | **1146** | 4.58 | 0.911 | 1244 | **1.98** | 0.924 | 1234 | 3.15 | 0.848 | 1230 | 6.34 |
| 18 | 0.910 | **860** | 4.54 | **0.976** | 893 | **2.51** | 0.820 | 942 | 3.79 | 0.841 | 941 | 7.13 |
| 19 | 0.976 | **618** | 3.32 | 0.978 | 695 | **1.82** | **0.980** | 1060 | 2.47 | 0.846 | 765 | 5.26 |
| 20 | **0.973** | **703** | 3.53 | 0.950 | 719 | **2.00** | 0.971 | 796 | 2.74 | 0.869 | 795 | 6.08 |
| 21 | 0.974 | **470** | 2.51 | 0.970 | 512 | **1.70** | **0.975** | 532 | 1.86 | 0.863 | 528 | 5.05 |
| 22 | 0.973 | **367** | 1.98 | **0.983** | 427 | **1.44** | 0.907 | 438 | 1.72 | 0.869 | 436 | 4.65 |

Fosmid pool-based sequencing provides very long fragments and is characterized by much higher ratio of the number of SNPs to the number of reads than the standard data sets generated by high-throughput sequencing platforms. We consider the fosmid sequence data for chromosomes of HapMap NA12878 and again take the trio-phased variant calls from the GATK resource bundle [15] as the true haplotype sequences. We compare the performance of AltHap to those of H-PoP, BP, and HapTree and report the results in Table 2. As can be seen from Table 2, AltHap achieves the best CPR and MEC score for most of the chromosomes and is faster than BP and HapTree. H-PoP is the fastest among the considered schemes but its speed comes at the cost of reduced accuracy. The second most accurate method is HapTree. However, it is significantly slower than other methods which makes it use more challenging in practice. Since HapTree could not finish assembling haplotype of the 6^th^ chromosome in 48 hours, that result is missing from the table.

**Table 2.** Performance comparison of AltHap, H-PoP, BP, and HapTree applied to the Fosmid data set. HapTree could not finish assembling haplotype of the 6^th^ chromosome in 48 hours.

| Chromosome | AltHap |  |  | H-PoP |  |  | BP |  |  | HapTree |  |  |
| --- | --- | --- | --- | --- | --- | --- | --- | --- | --- | --- | --- | --- |
|  | CPR | MEC | t(sec) | CPR | MEC | t(sec) | CPR | MEC | t(sec) | CPR | MEC | t(sec) |
| 1 | **0.955** | 9731 | 18.38 | 0.848 | 9845 | **2.127** | 0.876 | **9567** | 40.18 | 0.915 | 9676 | 6501 |
| 2 | **0.955** | 9589 | 38.89 | 0.904 | **9444** | **2.16** | 0.848 | 9698 | 42.90 | 0.923 | 9802 | 7196 |
| 3 | **0.917** | 7311 | 29.40 | **0.917** | **7182** | **1.79** | 0.847 | 7587 | 30.61 | 0.907 | 7705 | 4847 |
| 4 | **0.927** | **5508** | 26.69 | 0.926 | 5775 | **1.76** | 0.869 | 6288 | 31.10 | 0.908 | 6500 | 8392 |
| 5 | 0.920 | **6711** | 27.39 | **0.939** | 6910 | **1.95** | 0.863 | 6975 | 36.94 | 0.908 | 7094 | 5670 |
| 6 | **0.909** | **7213** | 33.68 | 0.885 | 7505 | **2.40** | 0.850 | 7590 | 41.20 | - | - | - |
| 7 | 0.907 | **6151** | 28.60 | **0.919** | 6829 | **1.68** | 0.858 | 6091 | 36.94 | 0.915 | 6169 | 5589 |
| 8 | **0.912** | **5927** | 23.82 | 0.902 | 6143 | **1.89** | 0.873 | 6282 | 38.87 | **0.912** | 6379 | 8316 |
| 9 | **0.918** | **5347** | 19.40 | **0.918** | 5719 | **1.76** | 0.851 | 5493 | 26.13 | 0.917 | 5513 | 4465 |
| 10 | 0.901 | **6044** | 24.07 | **0.924** | 6328 | **1.48** | 0.864 | 6503 | 27.65 | 0.889 | 6553 | 4838 |
| 11 | **0.908** | **5424** | 21.73 | 0.903 | 6432 | **1.68** | 0.858 | 5579 | 20.56 | 0.905 | 5625 | 5183 |
| 12 | **0.915** | **5456** | 24.25 | 0.914 | 5653 | **1.46** | 0.850 | 5706 | 24.19 | 0.913 | 5770 | 5654 |
| 13 | **0.904** | **3646** | 14.23 | 0.901 | 3708 | **1.54** | 0.827 | 3976 | 17.33 | 0.898 | 4029 | 5367 |
| 14 | **0.895** | 4156 | 18.64 | 0.891 | 4261 | **1.21** | 0.870 | 4004 | 14.84 | 0.906 | **4038** | 4103 |
| 15 | 0.900 | 4079 | 14.67 | 0.729 | **4001** | **1.06** | 0.823 | 4022 | 14.35 | **0.907** | 4116 | 3357 |
| 16 | 0.885 | 6197 | 26.28 | 0.715 | 6119 | **1.20** | 0.844 | **5112** | 29.51 | **0.942** | 5142 | 9683 |
| 17 | 0.897 | **4507** | 16.35 | 0.883 | 4911 | **1.22** | 0.876 | 4749 | 18.29 | **0.931** | 4806 | 3003 |
| 18 | **0.930** | **3080** | 12.68 | 0.908 | 3315 | **1.14** | 0.855 | 3457 | 13.31 | 0.919 | 3493 | 2303 |
| 19 | 0.857 | 4212 | 13.40 | 0.863 | 4115 | **0.84** | 0.835 | 3928 | 13.44 | **0.928** | **3953** | 1984 |
| 20 | **0.903** | **3512** | 13.64 | 0.900 | 4121 | **0.85** | 0.849 | 3814 | 15.97 | 0.901 | 3886 | 1529 |
| 21 | **0.927** | **1871** | 6.20 | 0.919 | 1974 | **0.68** | 0.872 | 1953 | 8.18 | 0.921 | 1979 | 1410 |
| 22 | 0.851 | 3654 | 17.24 | 0.878 | 3757 | **0.62** | 0.867 | 3910 | 14.72 | **0.924** | **3307** | 1351 |

### 5.2 Simulated data: the diploid case

To further benchmark performance of the proposed scheme, we test it on the synthetic data from [18] often used to compare methods for haplotype assembly of diploids. These data sets emulate haplotype assembly under varied coverage, sequencing error rates and haplotype block lengths. We constrain our study to the assembly of haplotype blocks having length *m* = 700 bp (the longest blocks in the data set). The results, averaged over 100 instances of the problem, are given in Table 3. As evident from this table, AltHap outperforms other algorithms for nearly all the combinations of data error rates and sequencing coverage and is also much faster than BP and HapTree while being slightly slower than H-PoP.

**Table 3.** Performance comparison of AltHap, H-PoP, BP, and HapTree on a simulated diploid data set from [18] with haplotype block length *m* = 700.

| Error rate | Coverage | AltHap |  |  | H-PoP |  |  | BP |  |  | HapTree |  |  |
| --- | --- | --- | --- | --- | --- | --- | --- | --- | --- | --- | --- | --- | --- |
|  |  | CPR | MEC | t(sec) | CPR | MEC | t(sec) | CPR | MEC | t(sec) | CPR | MEC | t(sec) |
| 0.1 | 5 | **0.996** | 477 | 0.043 | 0.993 | **402** | **0.012** | 0.867 | 698 | 1.421 | 0.886 | 491 | 2.13 |
| 0.1 | 8 | **0.999** | **759** | 0.128 | 0.998 | 780 | **0.035** | 0.872 | 861 | 4.627 | 0.884 | 767 | 3.82 |
| 0.1 | 10 | **0.999** | 954 | 0.404 | **0.999** | **903** | **0.109** | 0.873 | 1130 | 13.58 | 0.873 | 963 | 4.03 |
| 0.2 | 5 | **0.909** | **941** | 0.061 | 0.877 | 1021 | **0.027** | 0.812 | 953 | 2.671 | 0.762 | 988 | 9.36 |
| 0.2 | 8 | **0.981** | **1458** | 0.141 | 0.889 | 1532 | **0.098** | 0.861 | 1847 | 6.897 | 0.808 | 1562 | 6.69 |
| 0.2 | 10 | **0.991** | **1836** | 0.394 | 0.915 | 2023 | **0.201** | 0.867 | 2485 | 10.13 | 0.827 | 1943 | 4.20 |
| 0.3 | 5 | 0.607 | 1228 | 0.069 | 0.618 | 1331 | **0.041** | 0.537 | 1677 | 3.235 | **0.646** | **1170** | 10.21 |
| 0.3 | 8 | **0.677** | 2022 | 0.145 | 0.657 | 2250 | **0.098** | 0.572 | 2469 | 7.982 | 0.657 | **2021** | 6.17 |
| 0.3 | 10 | **0.750** | **2558** | 0.375 | 0.712 | 2979 | **0.217** | 0.596 | 3114 | 15.32 | 0.651 | 2597 | 5.74 |

### 5.3 Simulated data: the polyploid case

The performance of AltHap in applications to haplotype assembly for polyploids was tested using simulations; in particular, we studied how AltHap’s accuracy depends on coverage and sequencing error rate. The generated data sets consist of paired-end reads with long inserts that emulate the scenario where long connected haplotype blocks need to be assembled. We simulate sampling of the entire genome using paired-end reads and generate SNPs along the genome with probability 1 in 300. In other words, the distance between pairs of adjacent SNPs follows a geometric random variable with parameter 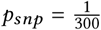 (the SNP rate). To emulate a sequencing process capable of facilitating reconstruction of long haplotype blocks, we randomly generate paired-end reads of length 500 with average insert length of 10,000 bp and the standard deviation of 10%; sequencing errors are inserted using realistic error profiles [12] and genotyping is performed using a Bayesian approach [15]. At such read and insert lengths, the generated haplotype blocks are nearly fully connected. Each experiment is repeated 10 times. AltHap is compared with H-PoP and BP. We also tried to run HapTree. However, HapTree could not finish the simulations for the considered block size in 48 hours.

Table 4 compares the CPR, MEC score, and running times of AltHap with those of H-PoP, and BP for biallelic triploid genomes with haplotype block lengths of *m* = 1000 for several combinations of sequencing coverage and data error rates. As can be seen there, AltHap outperforms both H-PoP and BP in terms of the CPR and MEC score in all the scenarios. In addition, AltHap is much faster than BP while capable of achieving highly accurate performance. Interestingly, AltHap is much faster than H-PoP for the highest coverage, i.e., 30. This is due to linear complexity of AltHap’s iterations which makes it suitable for high-throughput sequencing data characterized by high coverage. The results of tests conducted on simulated biallelic tetraploid genomes are summarized in Table 5, where we observe that AltHap outperforms the competing schemes in terms of both accuracy and running time.

**Table 4.** Performance comparison of AltHap, H-PoP and BP on a simulated biallelic triploid data set with haplotype block length *m* = 1000. HapTree could not finish the simulations in 48 hours.

| Error rate | Coverage | AltHap |  |  | H-PoP |  |  | BP |  |  |
| --- | --- | --- | --- | --- | --- | --- | --- | --- | --- | --- |
|  |  | CPR | MEC | t(sec) | CPR | MEC | t(sec) | CPR | MEC | t(sec) |
| 0.002 | 10 | **0.982** | **322** | 30.74 | 0.715 | 3642 | **14.17** | 0.689 | 4210 | 132.02 |
| 0.002 | 20 | **0.951** | **1986** | 59.65 | 0.731 | 7728 | **41.28** | 0.729 | 7762 | 416.44 |
| 0.002 | 30 | **0.984** | **2412** | **109.73** | 0.708 | 12865 | 265.22 | 0.697 | 14751 | 1310.18 |
| 0.01 | 10 | **0.917** | **1379** | 30.90 | 0.700 | 3786 | **14.18** | 0.681 | 4092 | 138.85 |
| 0.01 | 20 | **0.977** | **1597** | 60.09 | 0.709 | 8375 | **42.4** | 0.689 | 8601 | 460.60 |
| 0.01 | 30 | **0.989** | **3143** | **110.18** | 0.718 | 11769 | 266.39 | 0.681 | 15124 | 1301.54 |
| 0.05 | 10 | **0.971** | **2802** | 31.04 | 0.701 | 3978 | **14.53** | 0.669 | 4227 | 135.08 |
| 0.05 | 20 | **0.949** | **8222** | 59.95 | 0.703 | 9276 | **41.90** | 0.701 | 9484 | 460.49 |
| 0.05 | 30 | **0.826** | 17284 | **110.06** | 0.713 | **13778** | 268.05 | 0.676 | 16876 | 1315.84 |

**Table 5.** Performance comparison of AltHap, H-PoP and BP on a simulated biallelic tetraploid data set with haplotype block length *m* = 1000. HapTree could not finish the simulations in 48 hours.

| Error rate | Coverage | AltHap |  |  | H-PoP |  |  | BP |  |  |
| --- | --- | --- | --- | --- | --- | --- | --- | --- | --- | --- |
|  |  | CPR | MEC | t(sec) | CPR | MEC | t(sec) | CPR | MEC | t(sec) |
| 0.002 | 10 | **0.911** | **1113** | **43.56** | 0.707 | 3366 | 43.69 | 0.698 | 4568 | 290.29 |
| 0.002 | 20 | **0.950** | **2113** | **87.51** | 0.734 | 7359 | 113.77 | 0.712 | 9434 | 540.80 |
| 0.002 | 30 | **0.999** | **674** | **163.81** | 0.726 | 11693 | 598.39 | 0.715 | 12745 | 1496.51 |
| 0.01 | 10 | **0.982** | **938** | **44.66** | 0.693 | 3511 | 46.35 | 0.664 | 6475 | 296.27 |
| 0.01 | 20 | **0.993** | **1668** | **87.57** | 0.703 | 7882 | 114.86 | 0.669 | 10213 | 552.80 |
| 0.01 | 30 | **0.953** | **6518** | **164.71** | 0.710 | 12392 | 597.40 | 0.684 | 13245 | 1485.65 |
| 0.05 | 10 | **0.937** | **3905** | **44.96** | 0.677 | 4110 | 46.37 | 0.645 | 6869 | 306.29 |
| 0.05 | 20 | **0.958** | 9645 | **89.04** | 0.691 | **9109** | 118.96 | 0.685 | 11477 | 623.89 |
| 0.05 | 30 | **0.815** | 18690 | **165.04** | 0.700 | **14212** | 601.78 | 0.675 | 17681 | 1504.87 |

We further studied the performance of AltHap on triploid and tetraploid organisms having polyallelic sites and the results are summarized in Table 6 and Table 7, respectively. Notice that none of the competing schemes are capable of handling polyallelic genomes. The results suggest that AltHap was able to reconstruct underlying haplotype sequences with competitive performance at a low computational cost.

**Table 6.** Performance of AltHap on simulated polyallelic triploid data set with haplotype block length *m* = 1000. H-PoP, BP, and Hap-Tree cannot assemble polyallelic polyploid haplotypes.

| Error rate | Coverage | CPR | MEC | t(sec) |
| --- | --- | --- | --- | --- |
| 0.002 | 5 | 0.832 | 1377 | 43.05 |
| 0.002 | 10 | 0.932 | 897 | 115.13 |
| 0.002 | 15 | 0.935 | 1799 | 173.55 |
| 0.002 | 20 | 0.952 | 2346 | 232.07 |
| 0.01 | 5 | 0.747 | 2341 | 58.13 |
| 0.01 | 10 | 0.944 | 1269 | 115.41 |
| 0.01 | 15 | 0.909 | 3755 | 173.38 |
| 0.01 | 20 | 0.855 | 7272 | 235.86 |
| 0.05 | 5 | 0.799 | 3076 | 57.77 |
| 0.05 | 10 | 0.894 | 3925 | 116.33 |
| 0.05 | 15 | 0.931 | 6100 | 174.37 |
| 0.05 | 20 | 0.939 | 9120 | 236.73 |

**Table 7.** Performance of AltHap on simulated polyallelic tetraploid data set with haplotype block length *m* = 1000. H-PoP, BP, and Hap-Tree cannot assemble polyallelic polyploid haplotypes.

| Error rate | Coverage | CPR | MEC | t(sec) |
| --- | --- | --- | --- | --- |
| 0.002 | 5 | 0.794 | 2380 | 109.00 |
| 0.002 | 10 | 0.865 | 2043 | 220.6 |
| 0.002 | 15 | 0.938 | 2148 | 328.49 |
| 0.002 | 20 | 0.963 | 2388 | 432.28 |
| 0.01 | 5 | 0.797 | 2398 | 113.08 |
| 0.01 | 10 | 0.841 | 2927 | 220.33 |
| 0.01 | 15 | 0.828 | 5787 | 327.10 |
| 0.01 | 20 | 0.992 | 2319 | 432.85 |
| 0.05 | 5 | 0.746 | 4721 | 113.38 |
| 0.05 | 10 | 0.890 | 5146 | 211.43 |
| 0.05 | 15 | 0.923 | 7555 | 327.20 |
| 0.05 | 20 | 0.920 | 13704 | 435.15 |

The results of these extensive simulations imply that, as expected, haplotype assembly becomes more challenging as the number of haplotype sequences (i.e., the ploidy) increases. Nevertheless, in all the conducted studies, AltHap consistently reconstructs haplotype sequences accurately and with low computational cost. In addition, the results of Table 4 and Table 5 demonstrate that the computational time of AltHap grows significantly slower with coverage than the computational time of the competing schemes. In particular, for high coverages that are characteristic of high-throughput sequencing technologies, AltHap is the most efficient algorithm. Finally, we use the results obtained by running AltHap on simulated biallelic triploid data (i.e., the results summarized in Table 4) to examine tightness of the theoretical bounds on the CPR stated in Theorem 4.3. In particular, theoretical bounds on CPR are compared to the CPRs empirically computed for various combinations of coverage and data error rates (averaged over 10 independent problem instances). In Fig. 3a, the theoretical bound and experimental CPR results are shown as functions of the data error rate for coverage 15. We observe that the bound is reasonably close to the experimental results over the considered range of data error rates. In Fig. 3b, the theoretical bound and experimental CPR results are plotted against sequencing coverage for the data error rate *p*_*e*_ = 0.002. This figure, too, implies that the theoretical CPR bound is relatively close to the experimental results.

**Fig. 3.**
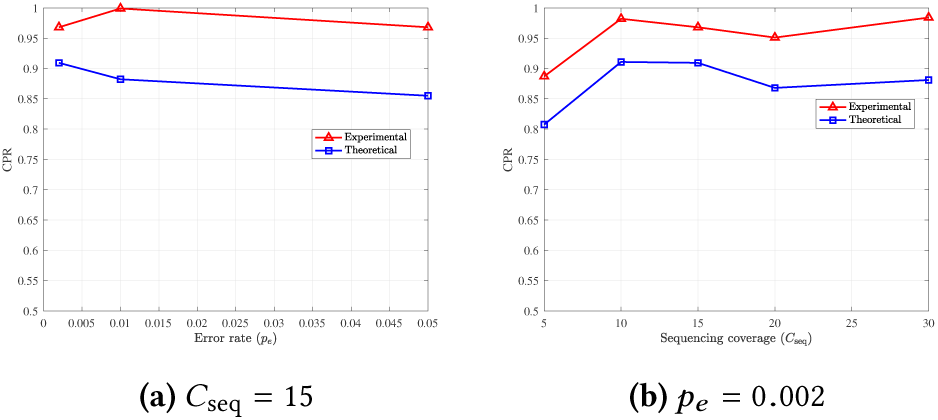
A comparison of the theoretical bound on CPR with the experimental results obtained by applying AltHap to the problem of reconstructing biallelic triploid haplotypes (synthetic data).

## 6 CONCLUSION

In this paper, we developed a novel haplotype assembly framework for both diploid and polyploid organisms that relies on sparse tensor decomposition. The proposed algorithm, referred to as AltHap, exploits structural properties of the problem to efficiently find tensor factors and thus assemble haplotypes in an iterative fashion, alternating between two computationally tractable optimization tasks. If the algorithm starts the iterations from an appropriately selected initial point, AltHap converges to a stationary point which is with high probability in close proximity of the solution that is optimal in the MEC sense. In addition, we analyzed the performance and convergence properties of AltHap and found bounds on its achievable MEC score and the correct phasing rate. AltHap, unlike the majority of existing methods for haplotype assembly for polyploids, is capable of reconstructing haplotypes with polyallelic sites, making it useful in a number of applications involving plant genomes. To the best of our knowledge, AltHap is the first polynomial time approximation algorithm for haplotype assembly with analytical guarantees on its achievable performance. Moreover, unlike several state-of-the-art techniques which are exponential in either read length or sequencing coverage, AltHap’s steps are linear in both and thus suitable to process sequencing data with long reads and deep coverage. Our extensive tests on real and simulated data demonstrate that AltHap compares favorably to competing methods in applications to haplotype assembly of diploids, and significantly outperforms existing techniques when applied to haplotype assembly of polyploids.

As part of the future work, it is of interest to extend the sparse tensor decomposition framework to viral quasispecies reconstruction and recovery of bacterial haplotypes from metagenomic data.

## APPENDIX Derivation of the proposed step size

For the proposed algorithm to converge, it must hold that

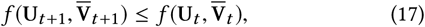

∀*t*. First, by noting (9) it holds that 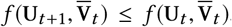. It now remains to show that 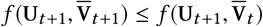. First, for the sake of notations and clarity, define

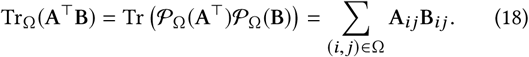

Recall that 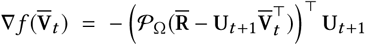 and 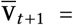 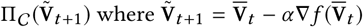. Since Π_*C*_ is a projection onto a convex set of constraints, *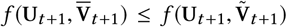*. Thus, it remains to show *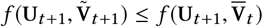*. Given that,

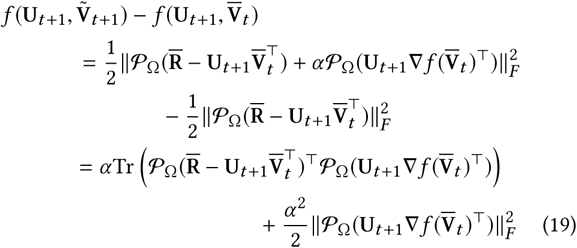

Now, consider the second term in the last line of (6). Following straightforward linear algebra we obtain

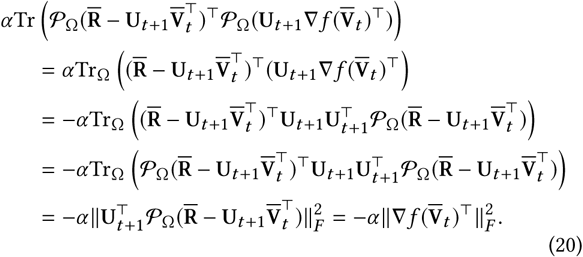

Therefore,

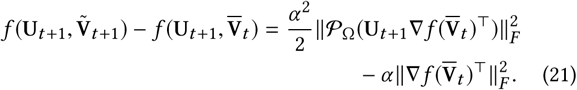

By choosing the step size (11) we obtain

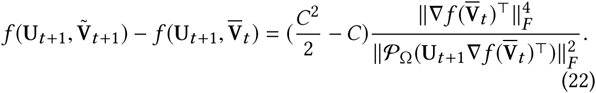

Clearly if *C* ∈ (0, 2) it must hold that *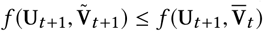*, which in turn implies convergence.

## Derivation of the MEC and CPR bounds

Recall that under conditions of Theorem 4.1, with probability 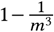 it holds that 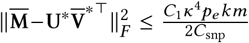. Once the stationary point 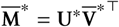 is found, Althap performs a decoding (rounding) step in order to obtain the binary solution 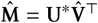. In this rounding procedure AltHap first normalizes all unfolded fibers such that sum of entries of each fiber equals 1. Then AltHap sets the largest entry of each unfolded fiber to 1 and the remaining three entries to 0. Eventually, AltHap reshapes the solution 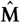 to the tensor 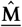. Note that this normalization is not required and we only consider it for the analysis purposes. Therefore, it is required to establish a bound on 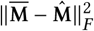. Let 𝓛 be the set of mismatching fibers, i.e., ∀*f* ∈ 𝓛, 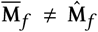. It is straightforward to see that 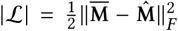. First, notice that 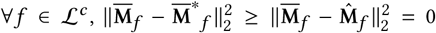.

In addition, consider a fiber ∀*f* ∈ 𝓛. The minimum value of 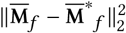 occurs when two entries in *f* are both equal to 0.5 and the remaining two entries are both 0. Hence, it becomes clear 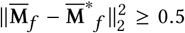. Thus, with probability 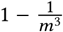 it holds that

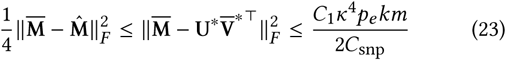

which is the desired relation. We now establish the MEC bound. Using the linearity of expectation and the above discussion, we obtain

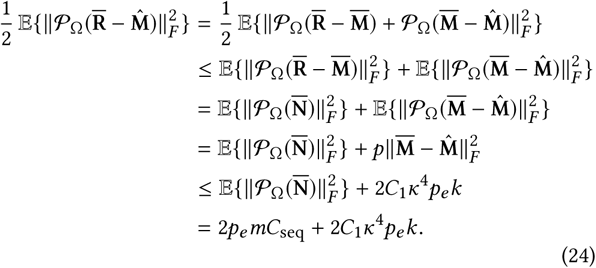

Thus, 𝔼{MEC} ≤ 2*p*_*e*_ (*C*_seq_*m* + *κ*^4^*C*_1_*k*). We now establish the CPR bound. The following is an equivalent definition of CPR computed using unfolded tensors of the true and the reconstructed haplotype sequences,

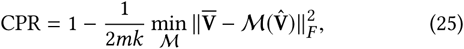

where 𝓜 is a one-to-one mapping from the corresponding entries of the lateral slices of 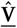 to those of 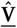. Assuming that sequencing reads uniformly sample haplotype sequences, on average, the mismatches between 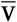 and 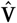 contribute equally to the number of mismatches between 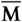 and 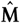. That is, 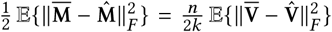. Therefore,

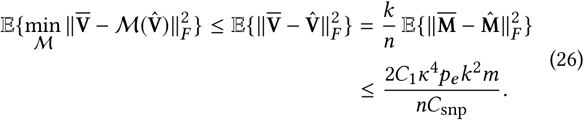

Thus, 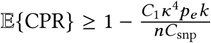 which is the desired bound.

## Footnotes

1 Notice that 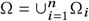.

2 For diploid data, we also quantified the performance of different methods by means of the switch error rate (SWER) (here omitted from brevity and reported at https://sourceforge.net/projects/althap/).

